# Wind drives drought responses of green roof vegetation in two substrates

**DOI:** 10.1101/486241

**Authors:** Arkadiusz Przybysz, Konstantin Sonkin, Arne Sæbø, Hans Martin Hanslin

**Affiliations:** Laboratory of Basic Research in Horticulture, Faculty of Horticulture, Biotechnology and Landscape Architecture, Warsaw University of Life Sciences – SGGW, Nowoursynowska 159, 02-776 Warsaw, Poland; Sagol School of Neuroscience, Tel Aviv University, 39040, Tel Aviv 6997801, Israel; Urban Greening and Environmental Engineering, The Norwegian Institute of Bioeconomy Research, Pb. 115, 1431 Ås, Norway

**Keywords:** green roofs, wind, drought survival, water balance, mortality, non-succulent species

## Abstract

The multifunctionality and delivery of ecosystem services from green roofs is improved by biological diversity of the roof vegetation. However, the frequency and intensity of drought episodes on extensive green roofs may limit the use of non-succulent species and the potential functional and phylogenetic diversity of the vegetation. Wind accelerates water use by plants and desiccation of the green roof substrate, and may be a key factor in selection of non-succulent plant species for green roofs. In this study, we tested wind interactions with green roof substrate composition and the effects on plant and substrate water balance, overall plant performance, and wilting and survival of three non-succulent species (*Plantago maritima* L., *Hieracium pilosella* L., and *Festuca rubra* L.) under realistic prolonged water deficit conditions. We found that, regardless of species or substrate tested, wind accelerated drought response. Drought-stressed plants exposed to wind wilted and died earlier, mostly due to more rapid desiccation of the growth substrate (critical substrate moisture content was 6-8%). The moderate wind levels applied did not affect plant performance when not combined with drought. Species with contrasting growth forms showed similar responses to treatments, but there were some species-specific responses. This highlights the importance of including wind to increase realism when evaluating drought exposure in non-succulent green roof vegetation.

## Introduction

Green roofs are engineered ecosystems representing an effective strategy to address some of the most challenging environmental issues in the urban areas [1, 2]. They provide several urban ecological goods and services, including thermal insulation to buildings, extension of roof lifespan, mitigation of the urban heat island, aesthetics, promotion of biodiversity (space, habitats and food) and stormwater management [1, 2]. This multifunctionality in delivery of ecosystem services by green roofs is improved by biological diversity of the roof vegetation [3]. Green roofs are however harsh environments, where the negative effects of drought, temperature extremes, radiation, air pollution, and wind on vegetation performance are amplified by the shallow and porous substrate low on organic matter used on extensive green roofs [2, 4, 5, 8]. The natural consequence of this green roof construction is limited water availability during prolonged drought, particularly when using water-demanding species [8, 9, 10]. Therefore, green roof vegetation for extensive roofs is usually selected based on species resistance to drought and heat in the root zone, resulting in prevalence of succulents, e.g., *Sedum* spp. [2, 7, 11, 12]. Succulents like *Sedum* and related *Crassulaceae* species fill however a rather narrow ecological niche. To increase the phylogenetic and ecological functionality of green roof vegetation, a wider range of species is required, including non-succulent species.

The most critical factor for long-term performance of non-succulent species is the intensity and frequency of drought episodes, even in cold humid regions [13]. The water balance of the substrate is determined by the input through precipitation, retention (storage) and loss through evapotranspiration. Retention is determined by the thickness of the substrate and the water holding capacity as affected by porosity and organic matter content, while evapotranspiration is affected by environmental factors such as irradiance, temperature and wind.

The impact of wind on plants is extremely variable and largely depends on its speed, duration, and frequency combined with plant characteristics [14, 15]. The effect of wind on green roof vegetation can include acceleration of substrate desiccation through evaporation, induction of mechanical and morphological responses, changes in leaf microclimate, and disturbance of leaf gas and heat exchange [15, 16, 17, 18, 19]. Despite the importance of wind for water loss and its effect on vegetation performance, there is a lack of knowledge about how wind affects green roof vegetation either directly or through effects on the water balance.

We sought to address this knowledge gap by examining how wind interacts with the water holding capacity of the substrate to affect water balance and performance of non-succulent vegetation under prolonged drought. We also examined how selected leaf chlorophyll *a* fluorescence parameters are related to desiccation and whether these parameters can be used to predict plant mortality on wind exposed green roofs.

## Materials and methods

### Experimental approach

To test the interactions between wind and substrate composition and their effects on drought response in green roof non-succulent species, we conducted a pot experiment under greenhouse conditions to monitor plant wilting symptoms and performance. Three species, representing different growth forms, were established in two different green roof substrates and exposed to simulated wind, while watering was withheld to mimic prolonged drought. The experimental design was a combination of two levels of wind (moderate wind or calm conditions), two levels of watering (withheld or control), and two different growth media (water-holding capacity 37% or 46%) for each of the three species, giving a total of 12 treatment combinations and 240 pots. Pot was the experimental unit, with 4 and 16 replicates per species and wind combination for control and drought treatments, respectively.

### Plant material and experimental conditions

The three species selected for the experiment (*Plantago maritima* L., *Hieracium pilosella* L., and *Festuca rubra* L.) are non-succulent candidates for green roofs representing different growth forms and phylogenetic lines with relatively high tolerance to abiotic stresses typical for green roofs conditions. Individual plants were established from large seedlings in pots (10 cm x 10 cm x 11 cm) containing two contrasting green roof substrates designed for this experiment, substrate A (more mineral, 36.7% water-holding capacity) or substrate B (more organic, 46.1% water-holding capacity). Substrate characteristics are given in Table 1. All plants were allowed to establish to greenhouse conditions for 6 weeks before the start of the experiment, during which time they were watered to approximately field capacity 2 times per week to achieve realistic drought acclimation before the experiment. The ambient light (58°N) was supplemented with additional 300 µmol m^−2^s^−1^ PAR (photosynthetically active radiation) daily from 7.30 to 19.30. Ventilation of the greenhouse was set to avoid air temperatures above 25°C.

**Table 1.** Composition (volume %) and characteristics of the two substrates used in the experiment.

| Parameter | Units | Substrate A | Substrate B |
| --- | --- | --- | --- |
| Compost | % | 5 | 10 |
| Shell sand | % | 5 | 0 |
| Pumice | % | 65 | 55 |
| Gravel | % | 25 | 25 |
| Biochar | % | 0 | 10 |
| pH (1:1, CaCl<sub>2</sub>) |  | 7.2 | 7.4 |
| Apparent density, dry | kg L<sup>-1</sup> | 0.84 | 0.71 |
| Max. water capacity | kg L<sup>-1</sup> | 0.37 | 0.46 |
| Total pore volume | % | 35.5 | 37.0 |

The greenhouse table was divided across into four parallel sections (1.0 m^2^) separated by 80 cm high Plexiglas plates. The two wind treatments were randomized on these sections and pots were randomized on drought treatment within sections. Pots were placed on a net frame on top of the greenhouse table, to increase substrate drainage.

### Drought and wind treatments

The day before water stress was initiated, all pots were watered to 100% field capacity and the weight of all individual pots was recorded. Drought stress was then imposed by completely withholding water for the drought treatment pots until the experiment was terminated after 30 days, when plants were harvested. Control plants were watered with 50 mL per pot every 2 days throughout the treatment period (corresponding to about 4 mm precipitation). Pot weight of drought-exposed plants chosen for measurements of chlorophyll *a* fluorescence and leaf temperature was monitored during the whole experiment by weighing, what allowed for estimation of gravimetric water content (GWC) based on the mass of water per unit mass of oven-dry soil determined after 48 h drying at 80^o^C at the end of experiment (Figure 1). The critical substrate moisture level, representing the threshold point of plant response, was also estimated.

**Fig 1.**
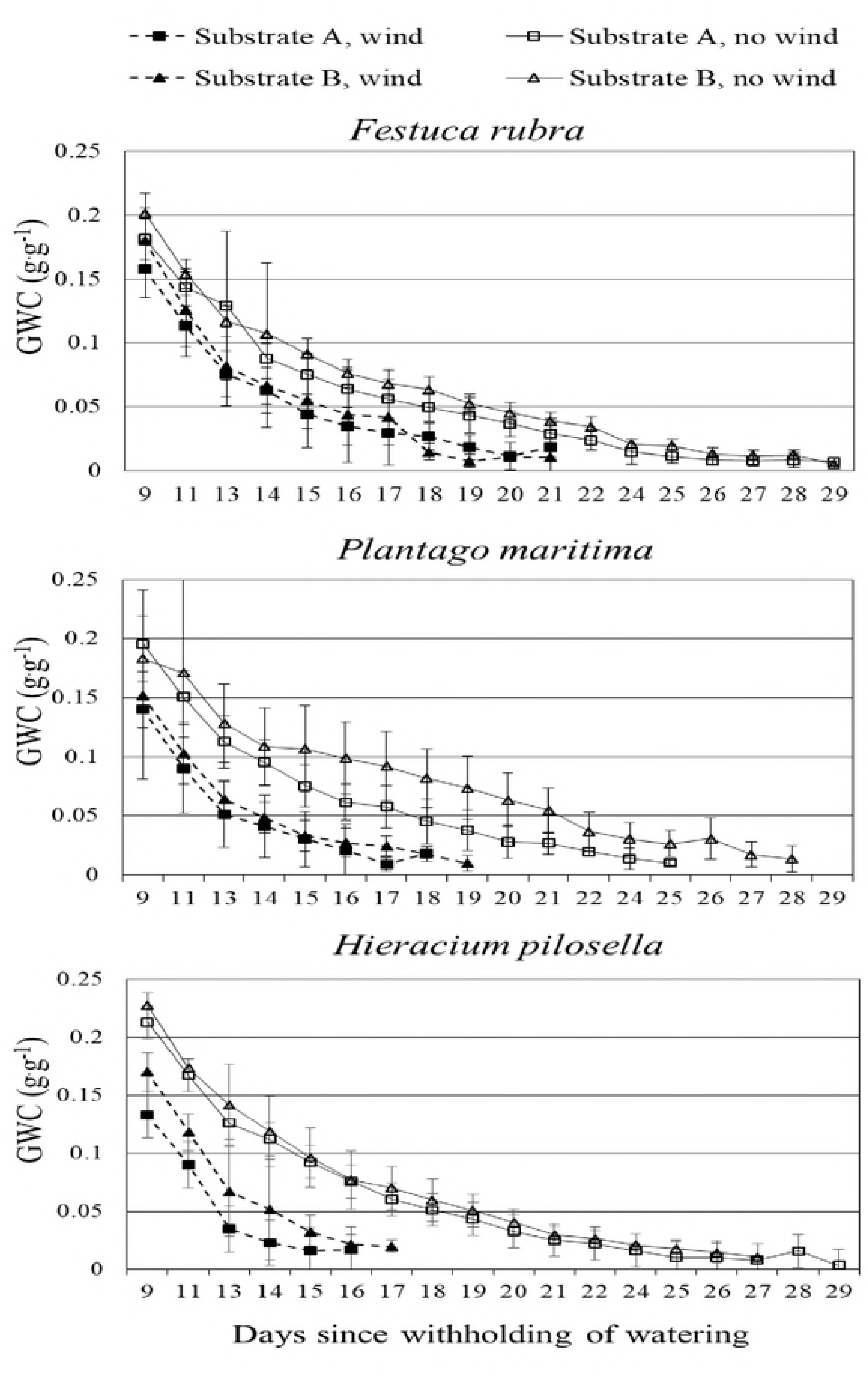
Effect of drought and wind on gravimetric water content (GWC) of green roof substrate under three non-succulent plant species. Values shown are mean with 95% confidence interval, n = 6.

Two of the parallel table sections received wind daily from two inline induct fans mounted side by side, about 100 cm from the pots with plants. This arrangement of fans simulated wind blowing daily for 12 hours, from 14.00 to 02.00, at an average speed of 3.7 m s^−1^ (± 0.63 SD, averaged over a grid of 60 pot positions per section) and ensured even wind speed within and between the two wind exposed sectors. Pots in all sections were re-positioned daily (always after performing observations and measurements) within sections to minimize edge and front effects of spatial wind patterns and other spatial patterns. Two sections were free from wind and serve as control for wind treatment.

### Observations and measurements

In the morning, wilting symptoms and survival were scored on all individuals, while fluorescence *a* parameters were recorded for a subset of the pots per treatment combination (4 for control and 6 for drought-stressed plants). These measurements were conducted on the same days when pots were weighed for GWC. At the beginning of the drought treatment, measurements were taken every second day. Later, when changes in plants performance were more rapid, measurements were performed daily. Performance of control plants was evaluated in the same way. Additionally, on selected days (14, 17, 18, 19, 20, 21, 22, 24 and 27 days after initiation of drought stress) leaf temperature was measured midday at 13.00. All measurements were performed between 8.00 and 14.00, when wind treatments were off.

Wilting was scored on every individual plant, on a scale from 1 to 9, where 1-3 means that plants were in good condition, 4-6 means that plants showed clear and increasing symptoms of wilting, and 7-9 means that plants were considered to be dying. Survival rate of plants was expressed as percentage of living plants and was calculated as: (p_l_/p_t_) x 100%, where p_l_ is number of living plants and p_t_ is total number of plants. Time taken to reach score class 8 was estimated for each treatment combination and analyzed in an ANOVA model with species, substrate, wind, and segment nested within wind as factors, using Minitab 17 (Minitab Inc., State College, PA, USA). The grading was tested by re-watering selected plants removed from the experiment (at score 9) and the results confirmed that a score of 9 gave 100% mortality. There was little difference in the size of individual plants within species at the start of the drought experiment, so variation in initial plant size was ignored.

Plants performance was determined in both control and stressed plants by measuring chlorophyll *a* fluorescence throughout the experiment, always at the same time of day (8.00-10.00), using a portable non-modulated fluorimeter Handy PEA or pulse-modulated fluorimeter FMS 2 (both Hansatech Instruments Ltd., Pentney, King’s Lynn, Norfolk, England). The Handy PEA device was used in first 22 days of drought stress treatments, while subsequent measurements were made with FMS 2. Before measurements, plants were pre-darkened for 45-60 min using clips supplied by the manufacturer. Chlorophyll *a* fluorescence induction transients were measured when leaves were exposed to a strong light pulse (3000 µmol photons m^−2^s^−1^). The fast fluorescence kinetics (F0 to FM) were recorded for 50 µs to 1 second. The measured data were analyzed by the JIP test [20]. Plant vitality was characterized by maximum quantum efficiency of photosystem II photochemistry (Fv/Fm) according to the equation: Fv/Fm = (Fm – Fo)/Fm. Measured data were also used for calculation of performance index (PI_abs_) [20], but only for the first 22 days of the experiment, as this parameter could not be measured with the FMS 2 device. For each species, wind treatment, and substrate type combination, chlorophyll *a* fluorescence was measured in four control (optimally watered) plants and six drought-stressed plants, equally divided between two table sections. Measurements were always performed on the same plants, which were randomly selected at the beginning of the experiment. Chlorophyll *a* fluorescence was measured until the number of surviving drought-stressed plants decreased to two.

Leaf temperature in *P. maritima* and *H. pilosella* plants was measured at 13.00 with a Raytek MX4 infrared thermometer (Fluke Process Instruments) on the same plants used for chlorophyll *a* fluorescence measurements. Accurate determination of leaf temperature was not possible for *F. rubra*, because of the leaf shape and fragility. To correct for differences in ambient conditions between days, leaf temperature values were standardized according to: (Tm–Tc)/Tc, where Tm is measured temperature and Tc is average temperature of control plants growing in the same substrate (no drought, no wind). Measurements were made in four control (optimal watered plants) and six drought-stressed plants, equally divided between two table sections.

Temporal changes in observed responses, measured parameters and differences between treatments were compared using 95% confidence intervals.

### Results

Overall, the three tested species showed similar responses to wind. Moderate wind increased the rate of water loss from the system (Figure 1) and increased midday leaf temperature (Figure 2). Wind also shortened the time until a decline of examined chlorophyll *a* fluorescence parameters (Figure 3 and 4) and accelerated rates of wilting and mortality of plants in the drought treatment in all species (Table 2). The effect of wind on wilting and the time until plants showed signs of wilting were not affected by substrate composition (Table 2). Regardless of species, wind did not affect the watered control plants (Figure 5).

**Table 2.** Effect of wind exposure and substrate composition on days until wilting (score 8) after withholding watering in three non-succulent species, analyzed in an ANOVA model. Wind treatment was applied in replicate greenhouse table sections, with sections nested within wind treatment in the statistical model. The proportion of variation (PoV) explained by each factor is shown. Total degrees of freedom (df) = 177. R^2^ (adj) = 69%.

| Factor | df | F | P | PoV (%) |
| --- | --- | --- | --- | --- |
| Species | 2 | 47.0 | 0.002 | 11 |
| Substrate | 1 | 1.46 | 0.349 | 0 |
| Wind | 1 | 769 | 0.001 | 53 |
| Spec*Sub | 2 | 7.85 | 0.040 | 3 |
| Spec*W | 2 | 6.02 | 0.061 | 1 |
| Sub*W | 1 | 0.05 | 0.850 | 0 |
| Section(W) | 2 | 0.61 | 0.732 | 0 |
| Spec*Sub*W | 2 | 0.74 | 0.532 | 0 |
| Spec*Section(W) | 4 | 0.66 | 0.651 | 0 |
| Sub*Section(W) | 2 | 0.99 | 0.446 | 0 |
| Spec*Sub*Section(W) | 4 | 0.99 | 0.414 | 1 |
| Residual | 154 |  |  | 31 |

**Fig 2.**
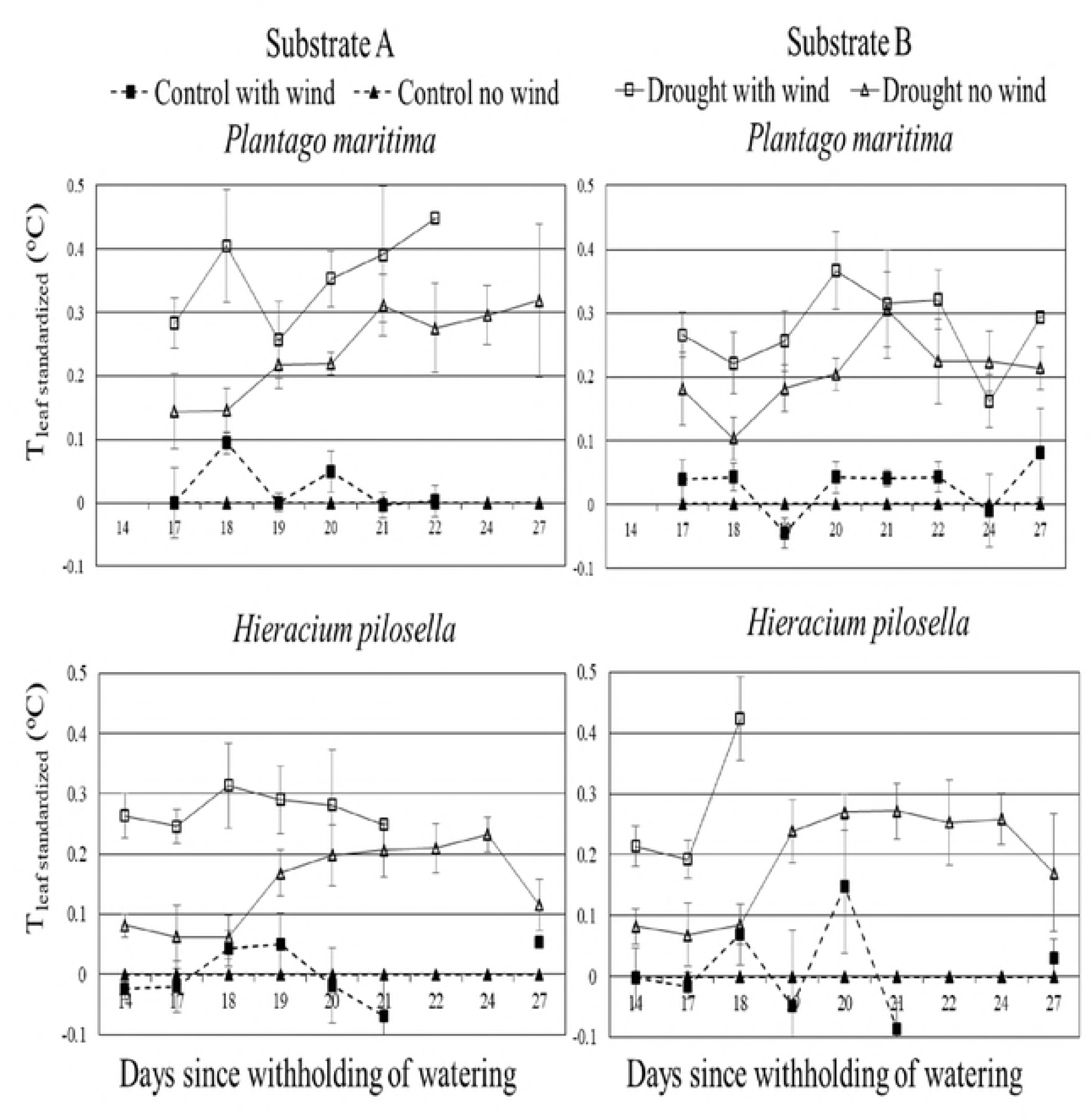
Effect of drought and wind on standardized midday leaf temperature in two candidate non-succulent green roof species. Values shown are mean with 95% confidence interval, n = 4 (control) or 6 (drought-stressed plants).

**Fig 3.**
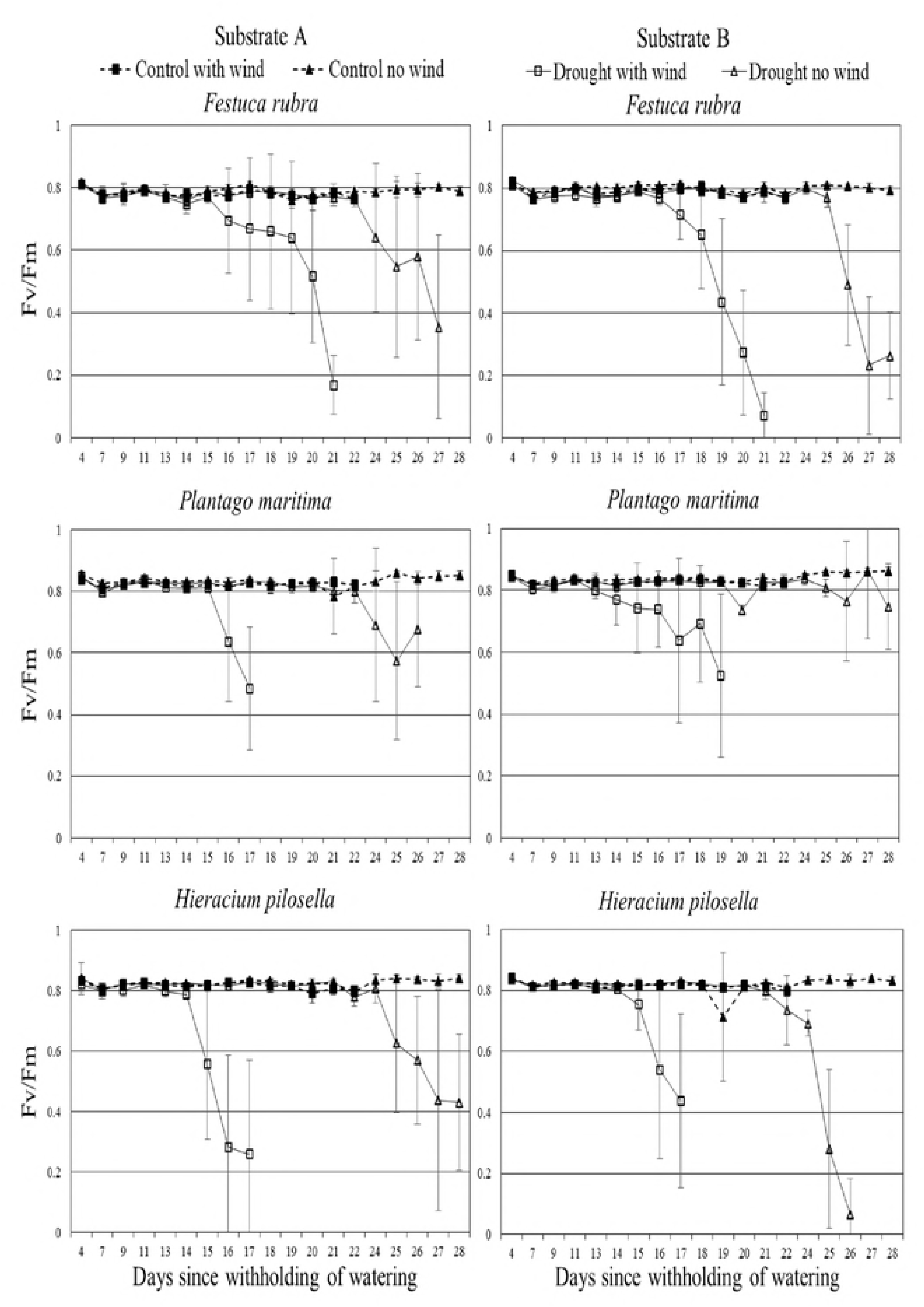
Effect of drought and wind on maximum quantum yield of photosystem II (Fv/Fm) of three candidate non-succulent green roof species. Values shown are mean with 95% confidence interval, n = 4 (control) or 6 (drought-stressed plants).

**Fig 4.**
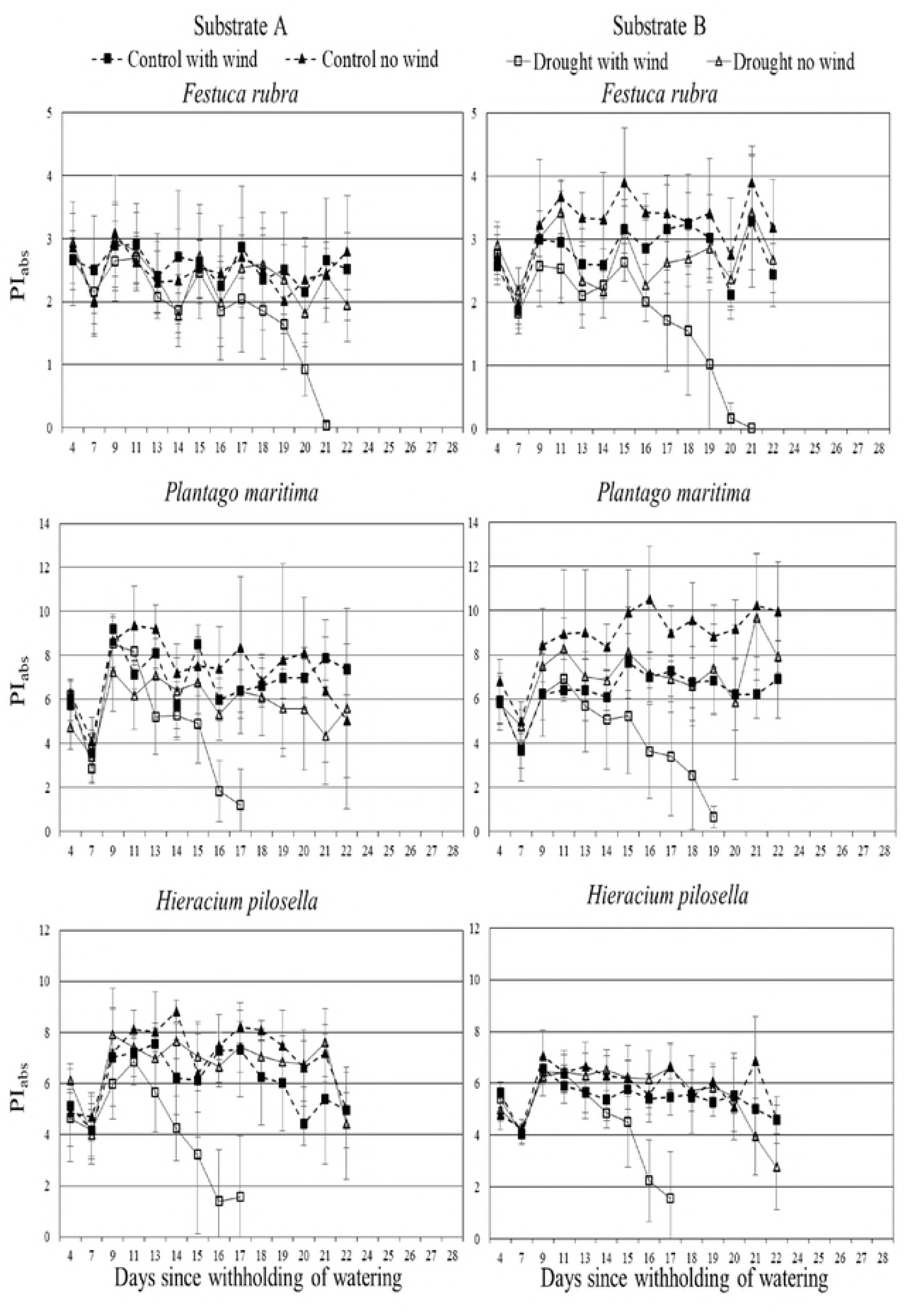
Effect of drought and wind on chlorophyll fluorescence performance index (PI) of three non-succulent species growing in two green roof substrates. Values shown are mean with 95% confidence interval, n = 4 (control) or 6 (drought-stressed plants).

**Fig 5.**
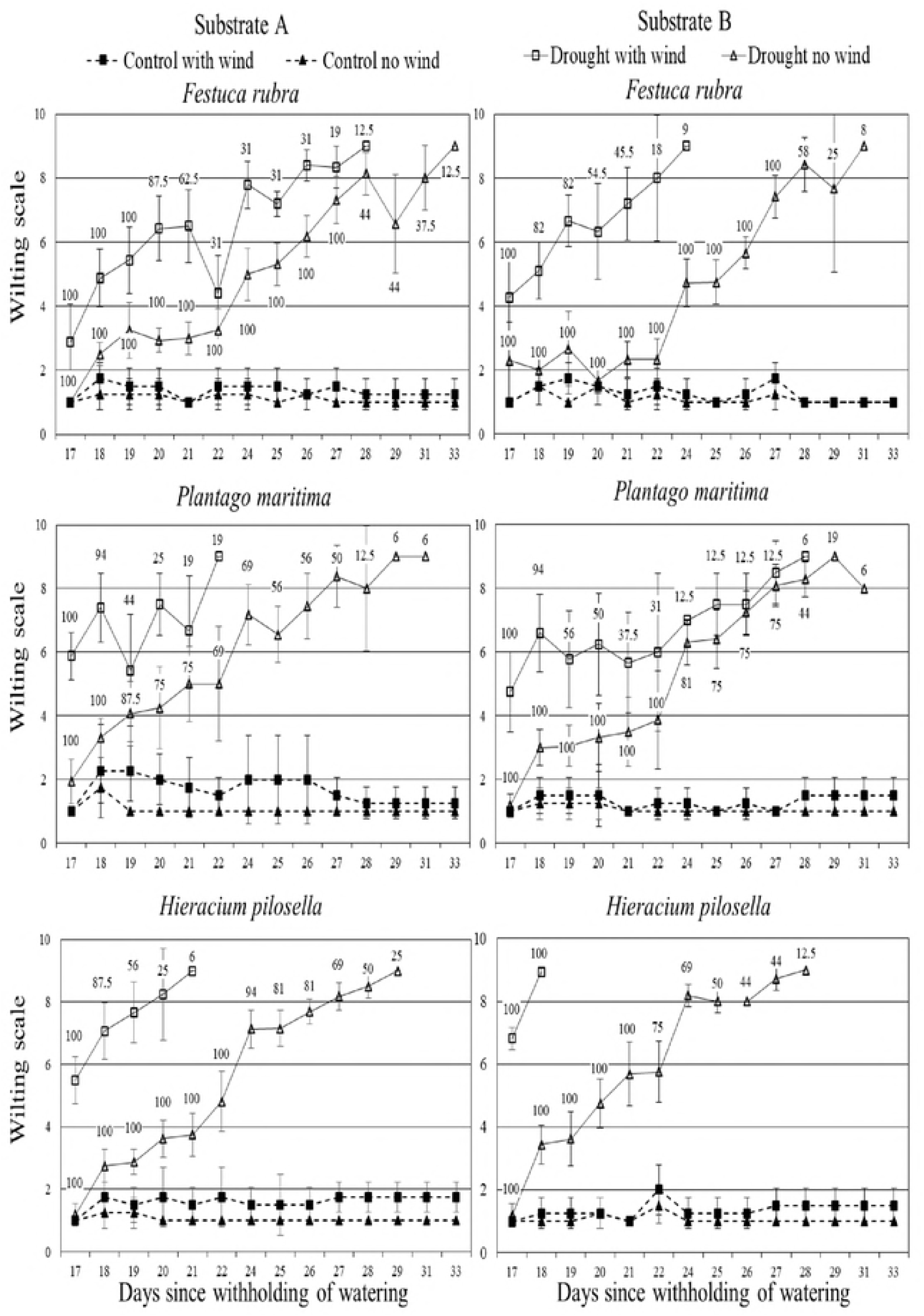
Effect of drought and wind on wilting and survival of three candidate species growing in two green roof substrates. Labels indicate percentage of living plants in each treatment, where 100% is initial number of plants. Values shown are mean with 95% confidence interval, n = 4 (control) or 6 (drought-stressed plants).

### Effects of drought

Due to different morphology and shoot biomass at the start of the drought period (*F. rubra* 230 ± 33 (SD) mg, *H. pilosella* 388 ± 66 (SD) mg, *P. maritima* 410 ± 49 (SD) mg), the three species had slightly different lags in response to drought, with *F. rubra* having a slower response and *H. pilosella* a faster response to negative pressure of water limited conditions (Figure 5). The three species also showed different strategies in their morphological responses to drought. *Plantago maritima* maintained turgor of all leaves until the rosette collapsed at the point of no return, *H. pilosella* maintained turgor of the youngest leaves with progressive wilting and curling of older leaves, while *F. rubra* maintained turgor of young leaves but gradually shed the oldest leaves. As expected, drought increased leaf temperatures (Figure 2) and decrease values of chlorophyll *a* parameters (Figure 3 and 4) over the experiment in all species.

### Effects of wind on willing and mortality under drought

Although responses to drought were species-specific, responses to wind were surprisingly similar between species, as no strong species by wind interactions were recorded (Table 2). Wind reduced the average time taken for the drought-stressed plants to reach stage 8 of wilting, by 4.7, 5.7, and 7.0 days in *P. maritima, H. pilosella*, and *F. rubra*, respectively (Figure 6). The first symptoms of wilting in drought-stressed plants (stage 4 on wilting scale) were also recorded sooner in wind-treated individuals, by three (substrate A) or eight (substrate B) days in *P. maritima*, six (substrate A) or four (substrate B) days in *H. pilosella*, and seven (substrate A) or eight (substrate B) days in *F. rubra* (Figure 5). The wind-induced acceleration of mortality of drought-stressed plants (plants considered dead and removed from experiment) was particularly evident in *F. rubra*, as the first plants of this species started to die nine (substrate A) or 10 (substrate B) days earlier when exposed to wind. *Plantago maritima* and *H. pilosella* maintained turgor for shorter periods than *F. rubra* under drought, and thus their mortality was accelerated by wind only by one (substrate A) to seven (substrate B) days and four (substrate B) to seven (substrate A) days, respectively. The significant species by substrate effect was due to a slightly better performance of *P. maritima* in the more organic substrate (B) and a slightly better performance of *P. maritima* and *H. pilosella* in the mineral substrate (A) (Figure 6, Table 2).

**Fig 6.**
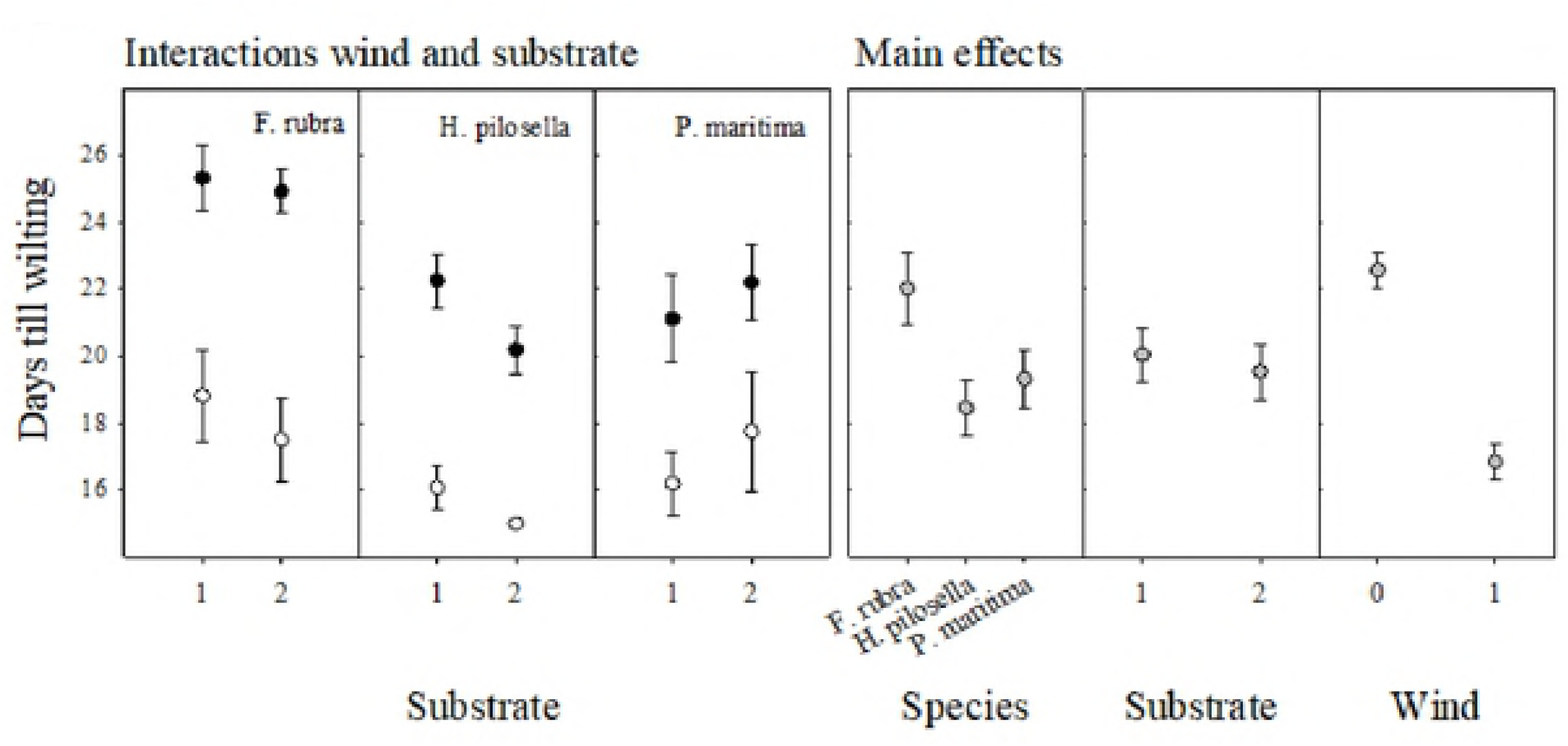
Average time until stage 8 (fatal) wilting in plants of three candidate green roof species exposed to wind (white symbols) or not (black symbols) after withholding of water to two green roof substrates (1 = substrate A, 2 = substrate B). Grey symbols show the main effects averaged over treatments. Values shown are mean with 95% confidence interval.

### Effect of wind on leaf temperature and chlorophyll *a* fluorescence

Treatment with wind resulted in increased midday leaf temperature in *P. maritima* and *H. pilosella* plants grown in conditions of prolonged drought (Figure 2). It was especially evident in *H. pilosella* plants, in case of which leaf temperature of drought stressed plant was only slightly higher than in optimally watered plants for first 19 days after initiation of drought, while leaf temperature of plants exposed to simultaneous drought and wind was already substantially higher (Figure 2).

The effects of wind on maximum quantum yield of photosystem II (Fv/Fm) showed a sharp threshold response (Figure 3), with the first decrease in this parameter recorded when the substrate moisture reached a critical level of 6-8% (calculated by weight) (Figure 1), similar for all species. Wind accelerated the negative effect of withholding water on Fv/Fm by on average 7-9 days across species and substrates (Figure 3). The performance index (PI_abs_) decreased in plants treated with simultaneous wind and drought, on nearly the same days as Fv/Fm (Figure 4). Fv/Fm showed a threshold drop at a wilt score of about 7, indicating that a wilt score of 8 was a reasonable indicator of integrated plant response to environmental conditions (Figure 7). This also reflects the fact that plants showed wilting symptoms for some days before Fv/Fm was affected and during that time photochemical conversion efficiency was probably not altered and photoinhibition did not take place. Based on Fv/Fm values and regrowth/survival after watering, we were quite conservative in mortality verdicts. Thus plants were apparently dead some time before they were scored as dead, probably soon after the first decrease in Fv/Fm was recorded.

**Fig 7.**
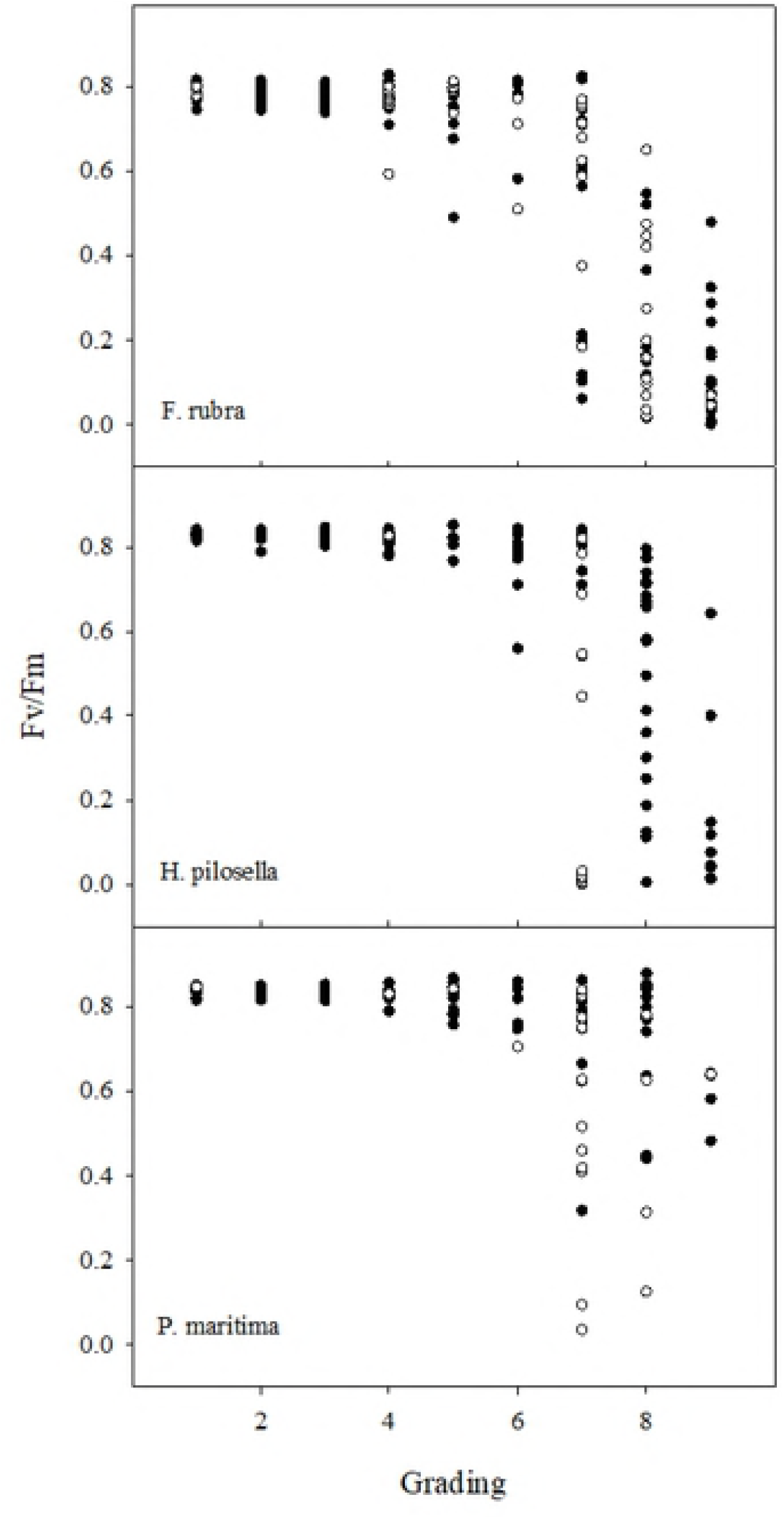
Relationship between wilting score and measured Fv/Fm under drought conditions for plants exposed to wind (white symbols) or not (black symbols) in the candidate green roof species *Festuca rubra, Hieracium pilosella*, and *Plantago maritima*.

## Discussion

### Effect of wind on drought responses of green roof vegetation

We found that wind had a strong negative effect on drought response in non-succulent candidate species for green roofs, and that this effect was not modified by the water-holding capacity of the two substrates used. We expected responses to wind to be slower for plants growing in the substrate holding more water, but this was not the case. Under water deficit conditions, wind increased the rate of water loss from the system, leading to higher midday leaf temperatures, a more rapid drop in the efficiency of the photosynthetic apparatus, and accelerated rates of wilting and mortality in the three species tested. These main effects of wind were in line with findings in other studies showing that wind may accelerate desiccation of green roof systems and, directly or indirectly, affect plant performance [15, 16, 17, 18, 19].

The effects of wind on plant performance were observed only when wind was combined with reduced water availability in the substrate. The moderate wind in our study (3.7 m s^−1^) was not sufficient to have an impact on the morphological and physiological parameters of watered control plants [21]. The effect of wind on plants depends on many factors, including its velocity and duration, but also on local environmental conditions, and can vary from positive to no or adverse impacts [15]. Therefore, it is possible that, under conditions different from those tested in this study; the response of well-watered plants to wind may be slightly different.

In general, wind was the factor that influenced drought-stressed plants the most, followed by differences between plant species, while substrate composition had no effect. Wind exposure accelerated wilting symptoms under drought conditions by 3-8 days and mortality by 1-10 days. The reason for the faster mortality of wind-exposed plants was increased water loss from both substrates. The critical substrate moisture level for all species and wind treatments was 6-8% (calculated by weight). Low water content and availability in the substrates most probably led to stomatal closure and decreased transpiration cooling, as indicated by the higher midday temperatures recorded in wind-exposed leaves, what was recorded throughout the drought period even if measurements were taken during periods when wind was turn off. The increased leaf temperature under abiotic stresses due to decreased gas exchange, particularly transpiration, is well known phenomenon [22]. In contrast, other studies have reported that wind (higher than 1.5 m s^−1^) usually reduces leaf temperature [23, 24] or at least brings it to the level of ambient temperature [25]. Leaf temperature higher than ambient has been reported under conditions of low wind (causing foliage heat transfer to decrease) and high radiation [23]. It was [19, 24, 26] that moderate wind, i.e., of similar strength to that used in this study, exacerbates the effects of drought not by stomata closing, but by increased transpiration rate due to reduced leaf boundary layers and increased vascular fraction. Boundary layer conductance increases linearly from approximately 10 mm s^−1^ at low wind speeds (< 0.1 m s^−1^) to over 150 mm s^−1^ at wind speeds of 2.0 m s^−1^ [23].

Wind exposure significantly accelerated the decline in Fv/Fm and PI_abs_ in drought-stressed plants. The PI_abs_ is a more responsive parameter to increased wind velocity (but also to other abiotic factors like salinity stress [27]) than Fv/Fm [28, 29]. However, in our study they showed very similar trends (at least for the first 22 days of drought when PI_abs_ was measured) and both remained at the optimal level when wind was applied to well-watered plants. Therefore, it appears that wind speed was within the tolerance range of the three species tested and that it exacerbated the symptoms of drought by drying the substrate, rather than directly affecting the photosynthetic apparatus. In fact, many wind-induced adaptive changes, on both molecular and morphological level, are associated with preventing soil and plant dehydration [30].

The three species tested showed similar main responses to wind under drought, although there were some differences in the time lag, with *F. rubra* and *P. maritima* showing a slower response than *H. pilosella*. This can be attributed to differences in plant size and to their morphology, as *F. rubra* has long narrow leaves, *H. pilosella* has a basal rosette with hairy leaves, and *P. maritima* has a basal rosette with semi-erect succulent leaves. All three species grow in habitats exposed to abiotic stresses; *H. pilosella* and the halophyte *P. maritima* grow on shallow soils, while *F. rubra* grows in a wider range of habitats. The Ellenberg soil moisture value, where 1 refers to plants preferring very dry conditions and 12 refers to submerged plants, is 5 for *F. rubra*, 4 for *H. pilosella*, and 7 for *P. maritima*.

We found the effect of wind to be independent of substrate composition (differences in content of organic matter and biochar) with the two substrates used. The water loss rates were primarily determined by wind and not substrate in all three species. This was surprising at first, as the substrates had water-holding capacity of 36.7% (substrate A) and 46.1% (substrate B), meaning that there was 0.1 kg more water per liter of substrate at field capacity in substrate B. Substrate B retained more water than substrate A over time, but the differences were small and not sufficient to affect the time until wilting. Of course, in other environmental conditions and for species better adapted to green roof conditions, increasing the water-holding capacity by addition of 10% organic matter and 10% biochar, as in substrate B, may be sufficientto increase plant performance during drought events [31]. On the other hand, although organic matter content improves substrate nutrition and water-holding capacity, decomposition over time can cause shrinkage, altering substrate water retention and air-filled porosity over time and dramatically reducing longevity [32].

### Practical implications for selection of green roof species

The provision of ecosystem services by green roofs may result from complex interactions between substrate type, substrate depth, and plant species, and these interactions probably lead to trade-offs between services [5]. However, substrate manipulations are often limited by the weight-loading capacity of the roof [6]. Plant selection is therefore a critical aspect of extensive green roof design. Limited water availability strongly reduces vitality and the ecosystem functionality of plants [2]. Plant response to drought is therefore a consideration in green roof species selection and usually results in prevalence of succulents [7, 8], due to the fact that prolonged drought significantly challenges survival of non-succulent species [7]. However, the adaptive strategies of succulents limit their contributions to multifunctionality compared with other species, e.g., their usefulness in stormwater management may be suboptimal. Species better suited for this purpose are able to keep rainfall retention of green roofs at a relatively high level, but green roof plants may experience >50 days of drought stress per year, sometimes reaching a point of no return and permanent wilting of non-succulent species [8]. The failure of most species, even from appropriate habitats, demonstrates the need to evaluate potential plants on green roofs under extreme climate conditions [7]. The novel findings in our study confirm that assessments of green roof plant survival under drought conditions must include wind effects, as otherwise the negative effect of water deficit/or drought periods may be underrated. There still is a potential to design substrates with high water holding capacity using water absorbing gels and similar, but our results indicated that for substrates with representative water holding capacities, this effect may be marginal.

Another objective of this study was to test whether selected leaf chlorophyll *a* fluorescence parameters can be used to predict plant mortality on windy exposed green roofs. We found that decreases in the value of Fv/Fm and PI_abs_ accompanied morphological changes recorded with the help of a wilting scale. Plant response to the combined action of drought and wind was very rapid, and increased wilting score and declining values of chlorophyll *a* fluorescence parameters coincided with serious, probably irreversible, changes in green roof plants.

## Conclusions

We found that wind accelerated the drought response of non-succulent candidate species for green roofs and this effect was not modified by higher substrate water-holding capacity (46% compared with 37%). Drought-stressed plants exposed to wind wilted and died faster, mostly due to more rapid desiccation of the substrate, while wind alone did not affect plant drought response. Species with contrasting growth forms generally showed similar responses to exposure to wind and drought. However, there were some species-specific responses, highlighting the importance of including wind in evaluation of non-succulent green roof vegetation.

## Acknowledgements

This study was funded by the Regional Research Fund for Western Norway (project 239039) and a strategic institutional program to the Norwegian Institute of Bioeconomy Research (NIBIO) through the Research Council of Norway (project 248349/F40).

Konstantin Sonkin would like to thank the Norwegian Institute of Bioeconomy Research and Confederation of Norwegian Enterprise (NHO) presidential program in Norway for funding his training.

We are grateful to Monika Małecka-Przybysz for laboratory support and for technical editing of the entire manuscript.

